# Coin-tossing by Kinesin-1 Head and Tail Binding Collectively Drives Microtubule Patterns

**DOI:** 10.1101/2025.08.18.670853

**Authors:** Jashaswi Basu, Kajal Singh, Anita Jannasch, Chaitanya A. Athale

**Affiliations:** Div. of Biology, IISER Pune, IISER Pune, Dr. Homi Bhabha Road, Pashan, Pune 411008, India; Center for Plant Molecular Biology (ZMBP), University of Tübingen, Tübingen, Germany

**Author notes:** Correspondence: Chaitanya A. Athale.

**Keywords:** microtubule, transport, kinesin-1, gliding assay, tail, bending, oscillations, patterns, *in vitro* reconstitution

## Abstract

Intracellular microtubule-based transport depends on an essential plus-ended molecular motor, kinesin-1. The N-terminal ATP-dependent head driving motility and a C-terminal cargo interacting tail, both bind microtubules. Previously, the interplay of both domains were hypothesized to play a role in collective microtubule sliding patterns, but the mechanisms remained unclear. Here, we show that full-length *Drosophila* kinesin-1 generates striking spatiotemporal patterns in gliding assays including bending, looping, and oscillations as well as stop-and-go motion of filaments. In presence of the motor domain alone these patterns are absent, but with the tail only construct, microtubules bind passively, with the tail acting as a static anchor. An equimolar mixture of these two domain constructs reproduces these spatio-temporal patterns both in terms of increased bending with length and velocity distributions. Our results support a simple ‘coin-toss’ model, where either head or tail bind microtubules with equal probability, leading to antagonistic interactions that drives emergent motility patterns and velocity distributions. Thus the kinesin tail plays an active mechanical role beyond regulation or cargo binding, modulating microtubule behavior through inter-domain competition.

## Introduction

Kinesins are a major class of microtubule-dependent, largely plus-end directed motor proteins, first identified for their roles in fast axonal transport in vertebrate neurons (Brady, 1985) and squid axons (Vale et al., 1985), as well as in mitotic spindle assembly (Scholey et al., 1985). Since then, many kinesin isoforms have been identified, with kinesin-1 being the best-characterized (Hirokawa et al., 2009). Kinesin-1 heavy chain (Khc) is a homodimer with an N-terminal ATP-dependent motor domain (Jon Kull et al., 1996), a C-terminal tail containing an ATP-independent microtubule binding site (Straube et al., 2006) and an auto-inhibitory motif (IAK) (Kaan et al., 2011). Kinesin-1 motors often act collectively to transport cargos (Shubeita et al., 2008).

*In vitro*, single kinesin-1 molecules walk processively along microtubules at velocities around 1,*µ*m/s (Yildiz et al., 2004; Visscher et al., 1999). In gliding assays with many motors, microtubule velocity remains largely unchanged across different motor densities (Howard et al., 1989). However, this behavior depends on the length of the kinesin stalk: shorter constructs like K401 slow down more at higher surface density compared to longer constructs such as K612 (Bieling et al., 2008; Crevenna et al., 2008). While these studies highlight motor and stalk contributions, the role of the tail domain in collective motility remains unclear.

The kinesin-1 tail is highly conserved (Navone et al., 1992) and contains a secondary microtubule-binding domain (MTBD), enabling full-length kinesin-1 to bind two microtubules simultaneously (Straube et al., 2006). This has been linked to microtubule bending and bundling in *Drosophila* S2 cells (Jolly et al., 2010) and fungal hyphae (Straube et al., 2006). Human kinesin-1 tail peptides bind microtubules electrostatically (Seeger and Rice, 2010), and have been implicated in processes such as ooplasmic streaming (Lu et al., 2016), axon outgrowth (Jolly et al., 2010), saltatory transport (Moua et al., 2011), and microtubule polymerization dynamics (Seeger and Rice, 2010). Despite these findings, how the kinesin-1 head and tail domains interact to shape microtubule dynamics remains poorly understood.

Here, we investigate the interplay between kinesin-1 head and tail domains in shaping microtubule gliding *in vitro*. Using gliding assays, we compare a full-length *Drosophila* kinesin-1 construct (K980) to a truncated motor-only construct (K401). Full-length kinesin induces distinct spatial (bending, looping, oscillations) and temporal (stop-and-go) motility patterns absent with K401-driven transport. By reconstituting separate head and tail domains in defined ratios, we show that these patterns arise from competition between motile heads and static tails, revealing a minimal mechanism for kinesin-1 driven spatial microtubule regulation.

## Results

### Full-length kinesin-1 induces spatial patterns in microtubule gliding

To investigate how tail-mediated binding affects microtubule motility and spatial organization in gliding assays, a full-length *Drosophila* kinesin-1 heavy chain construct (K980) was compared with a truncated kinesin-1 head construct (K401) (Fig. 1A, B). The K980 construct has all the domains of kinesin heavy chain: motor, stalk, coiled coil-1, hinge, coiled coil-2 and the tail with its ATP independent microtubule binding as well as the IAK auto-inhibitory domain. The K401 construct containing only the motor and stalk domains and includes an additional C-terminal BCCP tag. To reconstitute microtubule ‘gliding assays’, K980 was non-specific immobilized on treated glass cover-slips, allowing either the motor head or the tail domain to face and interact with microtubules. Other orientations such as the autoinhibited conformation or those where neither domain is accessible, are unlikely to support microtubule binding. In contrast, K401 was specifically immobilized via biotin-streptavidin interaction, ensuring a defined head-up orientation on the glass surface (Fig. 1C).

**Figure 1:**
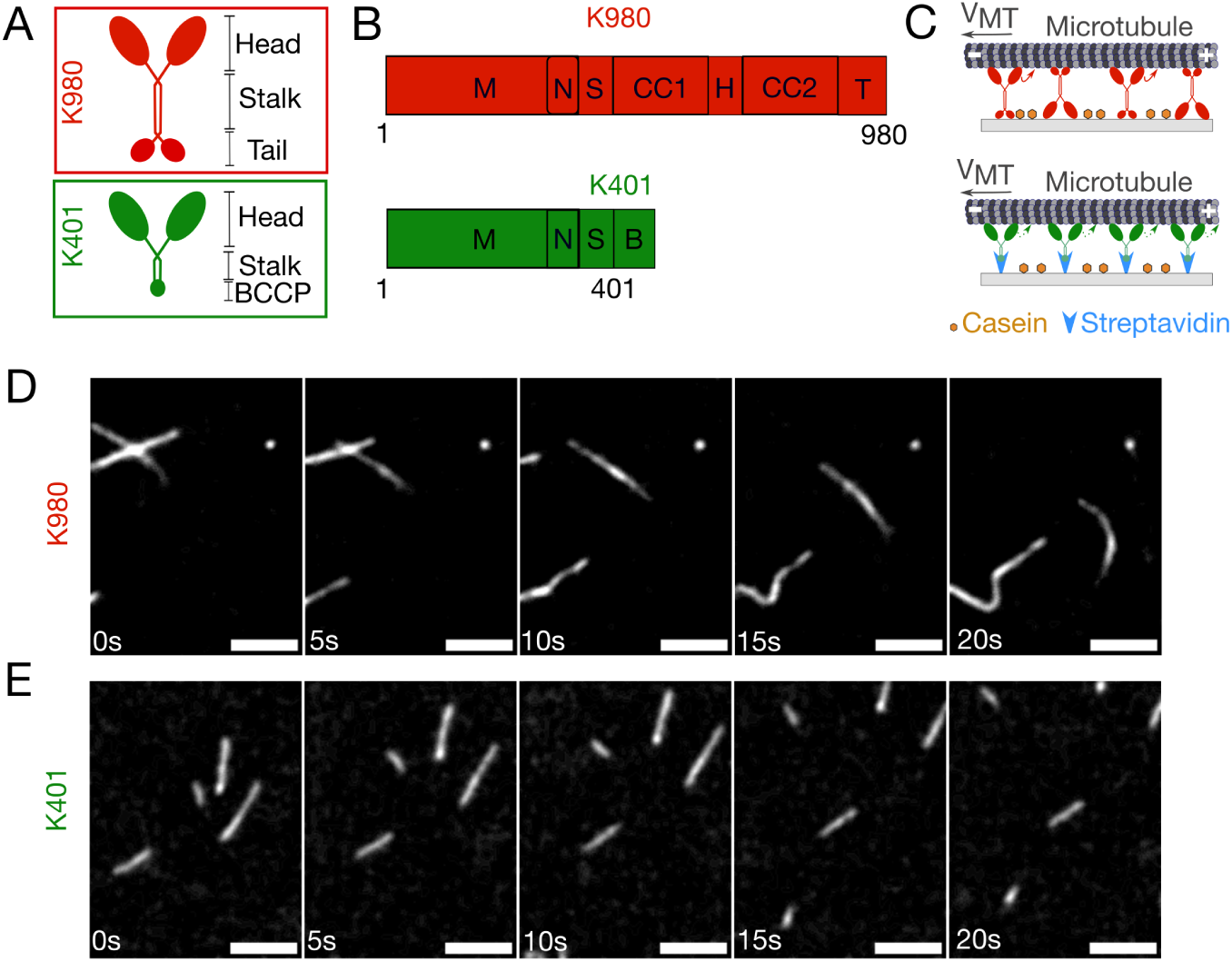
Full-length kinesin-1 and truncated head construct support microtubule gliding. (A) Schematic of full-length *Drosophila* kinesin-1 (K980, top) and truncated kinesin-1 head (K401, bottom) construct. (B) Domain structures of K980 (top) and K401 (bottom). M: motor domain with ATP-binding site; N: neck domain; S: swivel; CC1/CC2: coiled coils 1 and 2; H: hinge; T: tail; B: C-terminal BCCP tag present only in K401. (C) Schematic of the microtubule gliding assay showing K980 (top) and K401 (bottom) immobilized on a glass surface coated with casein. K401 binds via streptavidin for oriented attachment. Microtubules bind to surface-immobilized kinesin and glide with velocity *v*_MT_ (arrow). (D, E) Fluorescence time-series of rhodamine-labeled microtubules gliding on (D) K980 and (E) K401. Scale bar: 5 *µ*m.

In the presence of 1 mM ATP, both kinesin constructs supported active microtubule gliding. However, gliding assays with K980 frequently resulted in microtubules bending (Fig. 1D, Supplementary Video 1), in contrast to mostly straight microtubules observed in K401 driven gliding assays (Fig. 1E, Supplementary Video 2). We qualitatively classified K980-driven microtubule motility into three categories: (I) bending, (II) oscillations, and (III) looping (Fig. 2A, see Methods). To systematically quantify these patterns, the curvature of the microtubule was analyzed by the ratio of the length from end to end (*R*) to the contour length (L*_MT_*). This bending factor (*R/L*_MT_) estimates the degree of microtubule curvature (Fig. 2B; see Methods). A bending factor of ≈ 1 indicates a straight microtubule, while values *<* 1 reflect increasing curvature (Fig. 2B). The bending factor as a function of microtubule length showed distinct differences between gliding driven by K980 and K401. While K401 shows values ≈ 1, consistent with straight microtubules, K980 exhibits a clear length-dependent decrease in bending factor, indicating increased bending with microtubule length (Fig. 2C).

**Figure 2:**
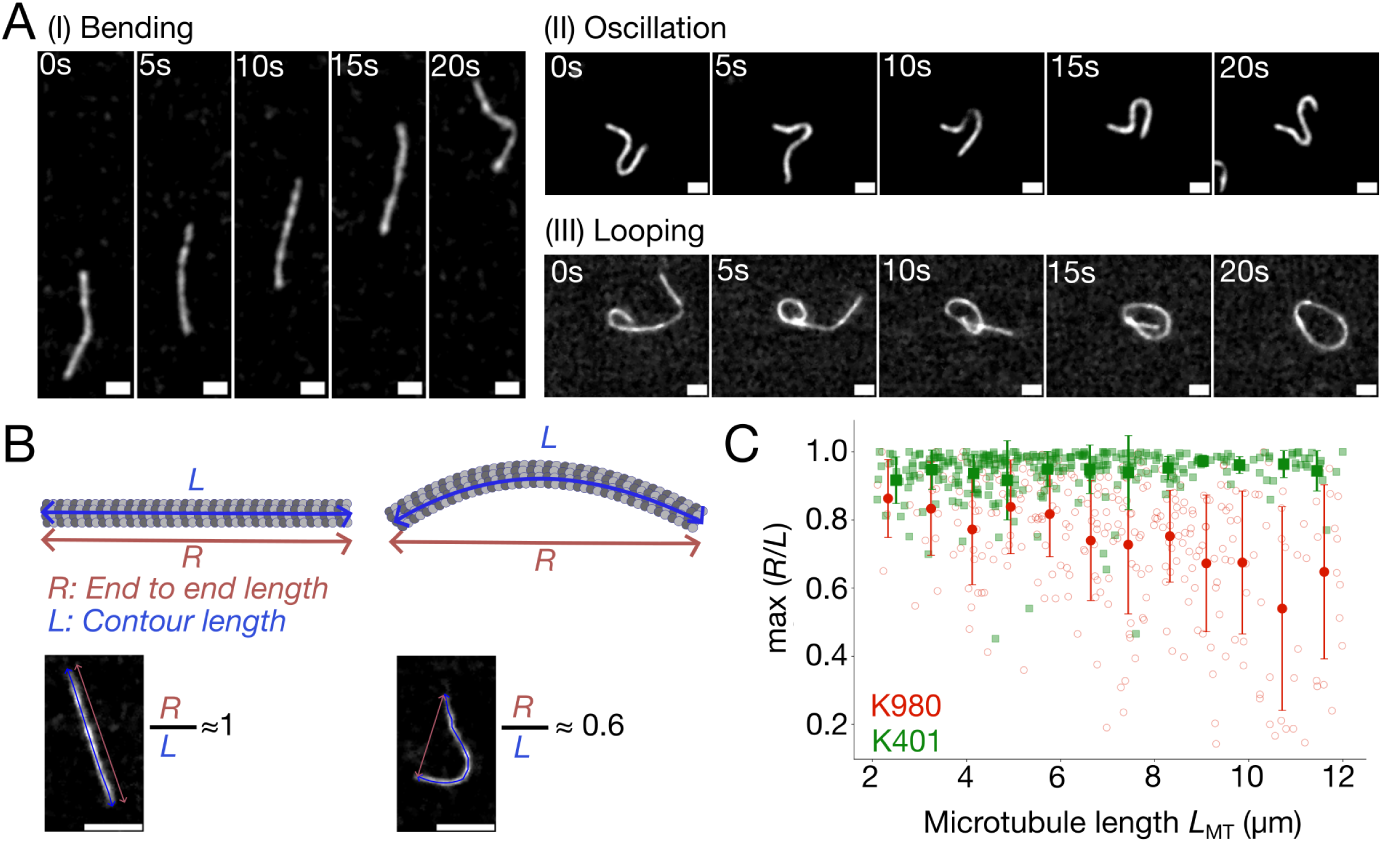
Full-length kinesin-1 transport leads to microtubule bending, unlike truncated kinesin head. (A) Time series of representative microtubules transported by K980 show (I) bending, (II) oscillations and (III) looping. Scale bar: 2 *µ*m. (B) (Top) Schematic showing microtubule end-to-end length (*R*, red) and contour length (*L*, blue). (Bottom) Two example microtubules are shown: one straight (*R/L* ≈ 1) and one bent (*R/L* ≈ 0.6). Scale bar: 4 *µ*m. (C) The maximal bending factor (*R/L*) plotted as a function of microtubule length (*L*_MT_) to compare microtubules transported by K980 (red) and K401 (green). Each data point represents the highest value (maximal) for an individual microtubule within a ∼20 s time interval. Large symbols indicate binned means±SD (bin width: 1 *µ*m, *N*_K980_ = 298, *N*_K401_ = 303).

Taken together, the striking occurrence of oscillations, loops, and a high pro-portion of bent microtubules suggests altered microtubule dynamics in full-length kinesin assays. These patterns were absent in assays using K401, which lacks the tail domain. To further investigate the underlying mechanism, we analyzed whether full-length kinesin-driven assays also exhibit interruptions in microtubule gliding motion.

### Full-length kinesin-1 causes stop-and-go microtubule gliding

Microtubule gliding dynamics were analyzed using kymographs and time-resolved trajectories of the leading microtubule tips (see Methods). For full-length kinesin K980 kymographs (Fig. 3A, top) and extracted trajectories (Fig. 3B, red lines) revealed three qualitatively distinct motility patterns: (a) processive, (b) stop-and-go, and (c) stop-and-bend. In contrast, microtubule gliding driven by the truncated kinesin head K401, showed mostly processive movement (Fig. 3A, bottom, Fig. 3B, green lines). To test whether this behavior depends on IAK-mediated autoinhibition, we used a mutant lacking the IAK motif (K936) (Supplementary Fig. S1A). Gliding assays with K936 (Supplementary Video 3) showed similar stop-and-go motion and a bimodal velocity distribution like K980 (Supplementary Fig. S1B), along with length dependent spatial patterns (Supplementary Fig. S1C). This suggests that the observed motility patterns are not due to autoinhibition.

**Figure 3:**
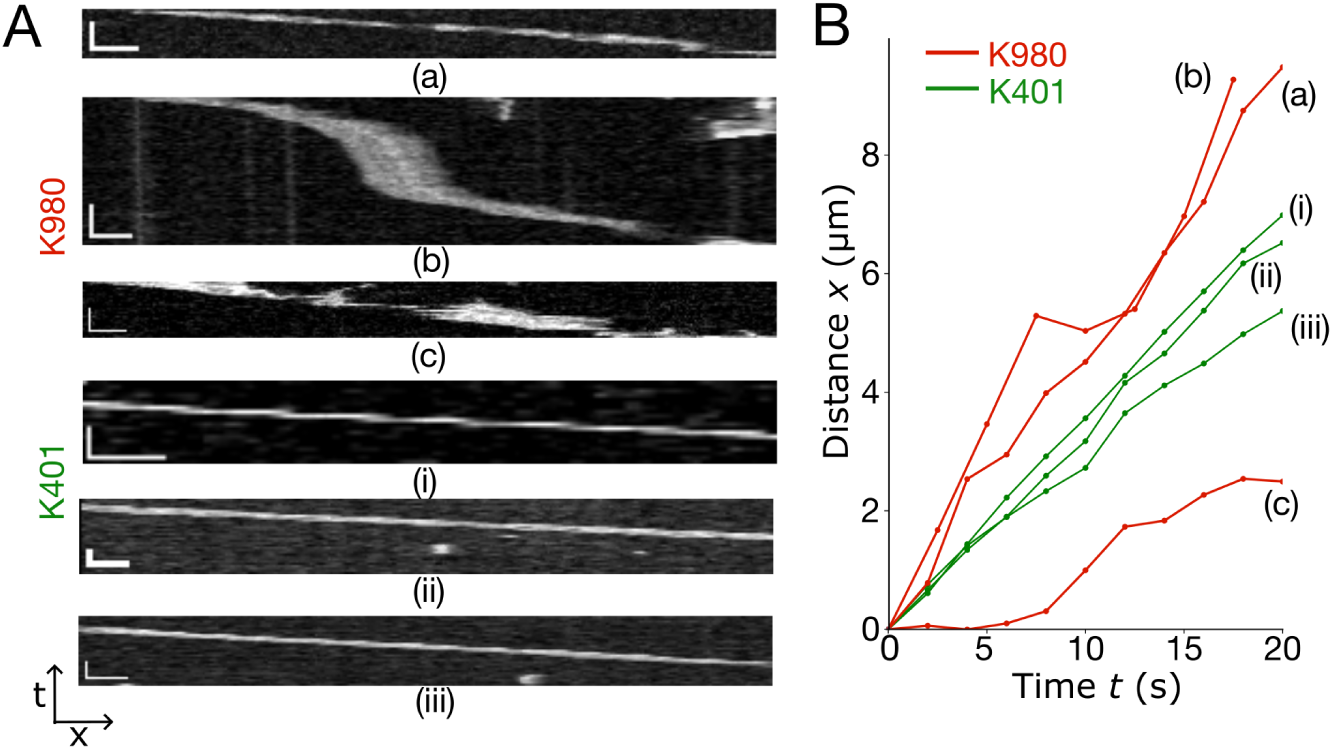
Full-length kinesin-1 induces stop-and-go microtubule transport. (A) Kymographs of individual microtubules driven by full-length kinesin-1 K980 (top) or truncated K401 (bottom). Vertical scale bar: 30 s; horizontal scale bar: 2 *µ*m. (B) Distance-time plots of microtubule leading ends for K980 (red) and K401 (green). K980-driven motion shows three modes: (a) processive, (b) stop-and-go, and (c) bending.

Thus we can show that full-length kinesin-1 collective transport results in stop-and-go microtubule transport, distinct from the processive gliding seen with the truncated kinesin head construct. To investigate the potential role of the tail domain in this behavior, we next characterized the microtubule-binding properties of the kinesin-1 tail domain.

### Kinesin-1 tail binds microtubules but does not support gliding

To evaluate whether the tail domain alone can account for microtubule binding, we purified a GFP-tagged construct comprising residues 910-980 of kinesin-1 (K910-980), which includes the reported microtubule-binding region and the IAK motif (Fig. 4A), based on previous reports (Lu et al., 2016; Winding et al., 2016). Microtubule binding was measured using a surface-immobilization assay, in which either the kinesin tail (K910-980) or the truncated kinesin head (K401) construct was anchored to the coverslip at equal concentrations (Fig. 4B). In both cases, microtubules bound to the surface, while no binding was observed in GFP-only controls (Fig. 4B). Quantification of microtubule surface density revealed no significant difference between K910-980 and K401 constructs (Fig. 4C), indicating that the minimal kinesin tail fragment binds microtubules with comparable efficiency to the kinesin head domain under these conditions. To confirm that binding alone does not imply motility, we tested whether the kinesin tail can drive microtubule transport (Supplementary Fig. S2A). As expected, the tail construct did not support microtubule gliding, with velocities near zero (Supplementary Fig. S2B,C, Supplementary Video 4). As a control, we also quantified the gliding velocity of a rigor mutant of full-length kinesin (K980(R210A)), which carries an R210A substitution in the motor domain that abolishes ATP hydrolysis (Farrell et al., 2002) and is known to lack motility (Mickolajczyk et al., 2015). This construct showed a similar gliding velocities near zero (Supplementary Fig. S2D).

**Figure 4:**
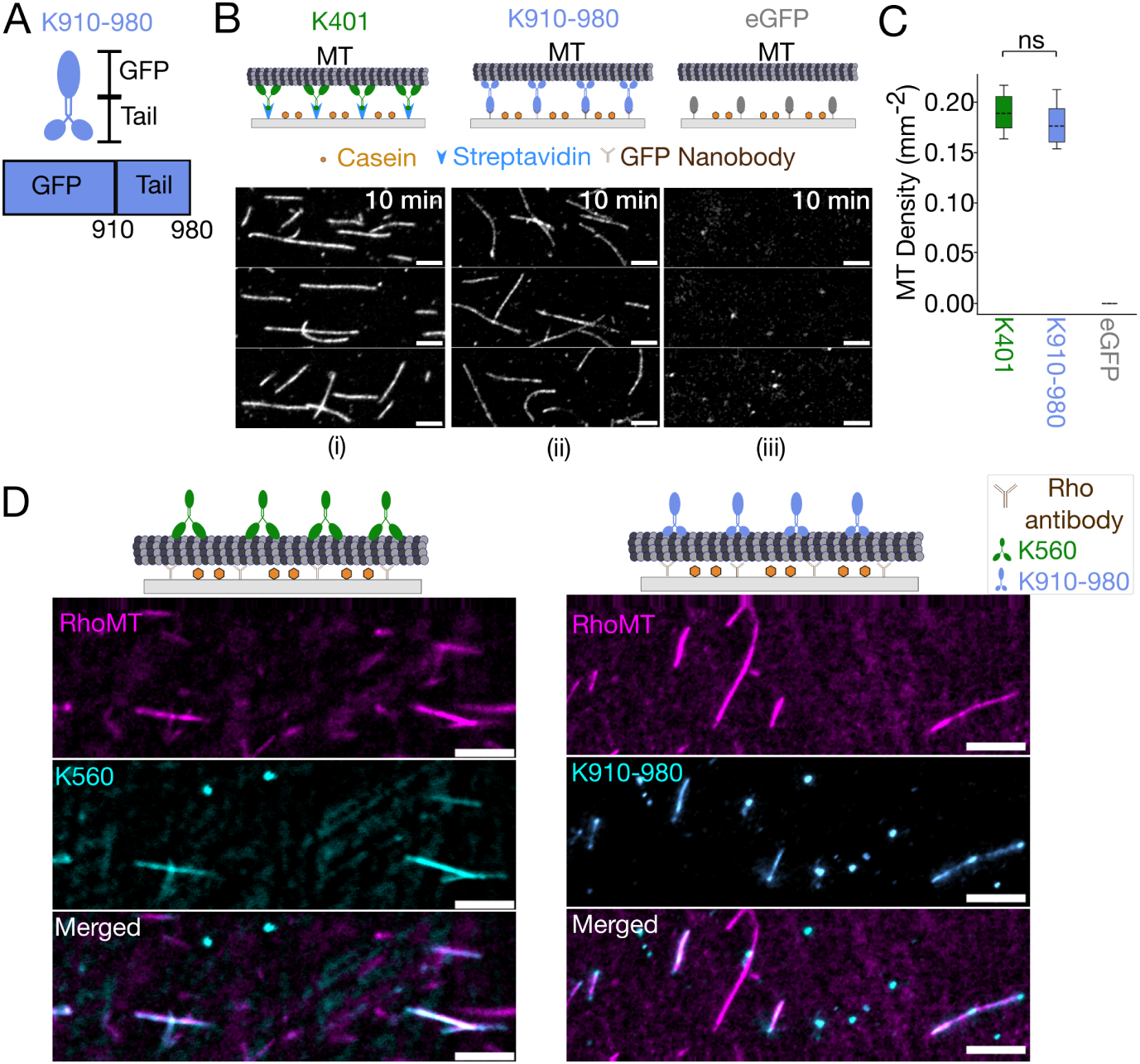
Kinesin-1 head and tail domains bind microtubules. (A) Schematic of the GFP-tagged kinesin-1 tail domain (K910-980). (B) Microtubules immobilized on coverslips coated with either (i) K401, (ii) K910-980 or (iii) eGFP. Scale bar: 2 *µ*m. (C) Microtubule density of microtubules immobilized on either K401,K910-980, or GFP (*N*_K401_ = 425, *N*_K9100-980_ = 416). ns: not statistical significant (D) GFP-tagged truncated kinesin head (K560, left) or tail domain (K910-980, right) were added to immobilized microtubules. Co-localization is shown in merged images. Scale bar: 3 *µ*m.

We further compared the microtubule-binding activity of the truncated kinesin head and the kinesin tail by adding either K560 (a truncated motor construct comprising residues 1-560, used here for its GFP tag) or K910-980 to surface-immobilized microtubules (Fig. 4D). Both constructs co-localized with microtubules, but K560 bound continuously along their length, whereas K910-980 was enriched at discrete regions.

Although microtubule binding by the kinesin tail has been reported *in vitro* (Straube et al., 2006; Seeger and Rice, 2010), its role in motility remains unclear. To assess whether tail binding influences motility when combined with motor activity, we reconstituted equimolar mixtures of kinesin head and tail domains and analyzed microtubule dynamics in this minimal system.

### Equal head-tail ratio reproduce kinesin full-length microtubule gliding behavior

In order to test whether the interplay between kinesin motor head and tail domains leads to the spatial motility patterns observed with full-length kinesin (K980), we reconstituted gliding assays using an equimolar mix of the truncated kinesin head (K401) and tail domain (K910-980). Both constructs were specifically immobilized on glass surfaces using streptavidin and GFP nanobody, respectively (Fig. 5A).

**Figure 5:**
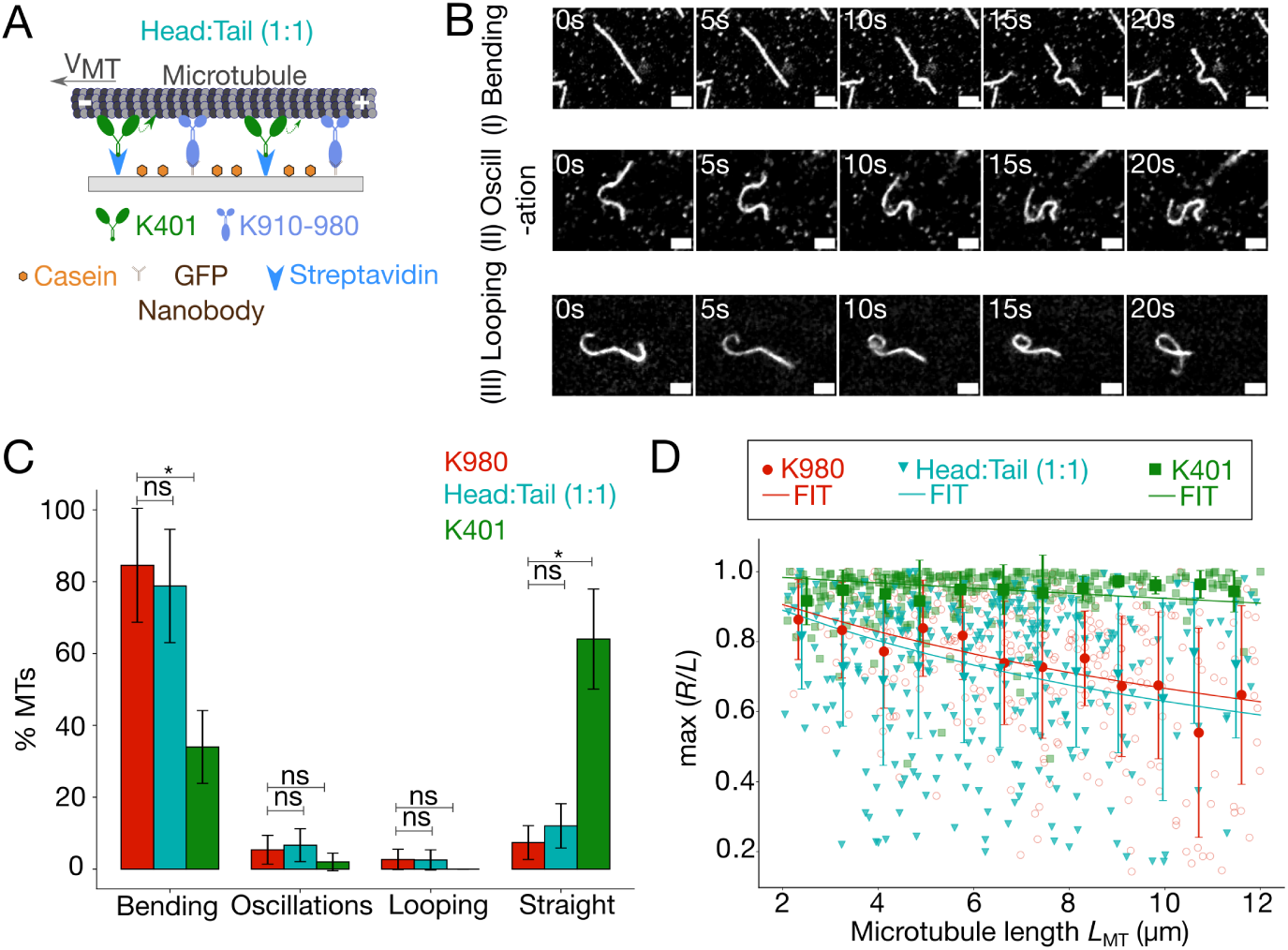
Equal head-tail mix reproduce spatial motility patterns of full-length kinesin-1. (A) Schematic of gliding assay with equimolar amounts of truncated kinesin head (K401) and tail (K910-980) immobilized via streptavidin and GFP nanobody, respectively. (B) Time series of distinct spatial microtubule motion patterns during gliding in the mixed head-tail assay: (i) bending, (ii) oscillation, and (iii) looping. Scale bar: 2 *µ*m. (C) Percentage of microtubules showing bending, oscillations, looping and straight motion for full-length kinesin (K980, red), mixed head-tail assay (cyan) and truncated head (K401, green). (D) The maximal bending factor (*R/L*) is plotted as a function of microtubule length (*L*_MT_) for microtubules transported by K980 (red), head-tail mix (dark cyan), and K401 (green). As before, each data point is the maximal value for an individual microtubule in ∼20 s. Large symbols indicate binned means±SD (bin width: 1 *µ*m). Dashed lines represent fits of binned averaged *R/L* to the Kratky-Porod model. Bending stiffness = 0.3121584 *µ*m^1^ (K980), 0.0495404 *µ*m^1^ (K401), 0.37231774 *µ*m^1^ (Head-Tail). (*N*_K980_ = 298, *N*_K401_ = 303, *N*_Head-Tail_ = 316)

Microtubule gliding in the reconstituted kinesin head-tail mix also exhibited dynamic behaviors, including bending, oscillation, and looping, similar to those observed with K980 (Fig. 5B; Supplementary videos 5, 6A, B). The distribution of these patterns was similar, with bent microtubules dominating in both cases (Fig. 5C).

To quantify microtubule curvature, we measured the bending factor (*R/L*) as a function of microtubule length. The kinesin head-tail mix showed a clear length-dependent decrease in the bending factor, matching the behavior observed with K980 (Fig. 5C). Mean bending factor values for the head-tail mix and K980 were not statistically different (Fig. 7C). K401-driven gliding have a bending factor of ≈ 1, consistent with straight microtubule trajectories.

The bending factor decreases with microtubule length as microtubules behave as semiflexible polymers and can be described with the Kratky-Porod worm-like chain model (Kratky and Porod, 1949; Mizushima-Sugano et al., 1983) (see Methods Eq. (1)). The fitting parameter, the bending stiffness *λ*, is proportional to the persistence length and reflects microtubule flexural rigidity, where larger *λ* denotes higher flexibility(Mizushima-Sugano et al., 1983; Hsu et al., 2010). To quantify microtubule flexibility, we fit the bin-averaged bending factor with respect to microtubule length for K980, K401 and head-tail mix (Fig. 5D).

Thus, an equimolar mixture of truncated kinesin head (K401) and kinesin tail K910-980 reproduces the spatial motility patterns observed with full-length kinesin (K980), including microtubule bending, oscillations, and looping. These results suggest that the spatial microtubule gliding patterns observed with K980 are primarily driven by additional microtubule binding through the inactive tail domain. To support this conclusion, we next test if the tail domain K910-980 also influences microtubule gliding velocity when mixed with K401.

### Equal head-tail ratio induces bimodal velocity distributions in microtubule gliding

We hypothesized that in full-length kinesin (K980) gliding assays, where kinesin is randomly immobilized, both the motor head and tail domains have equal chances of binding to microtubules-the ‘coin-toss’ model. This model would then predict multiple motor configurations bound to microtubules: head, tail, or a combined head-tail binding (Fig. 6A). Together, these states could explain the spatial and temporal motility patterns observed in K980 assays, as well as those reproduced by the equimolar head-tail mixture described in the previous section.

**Figure 6:**
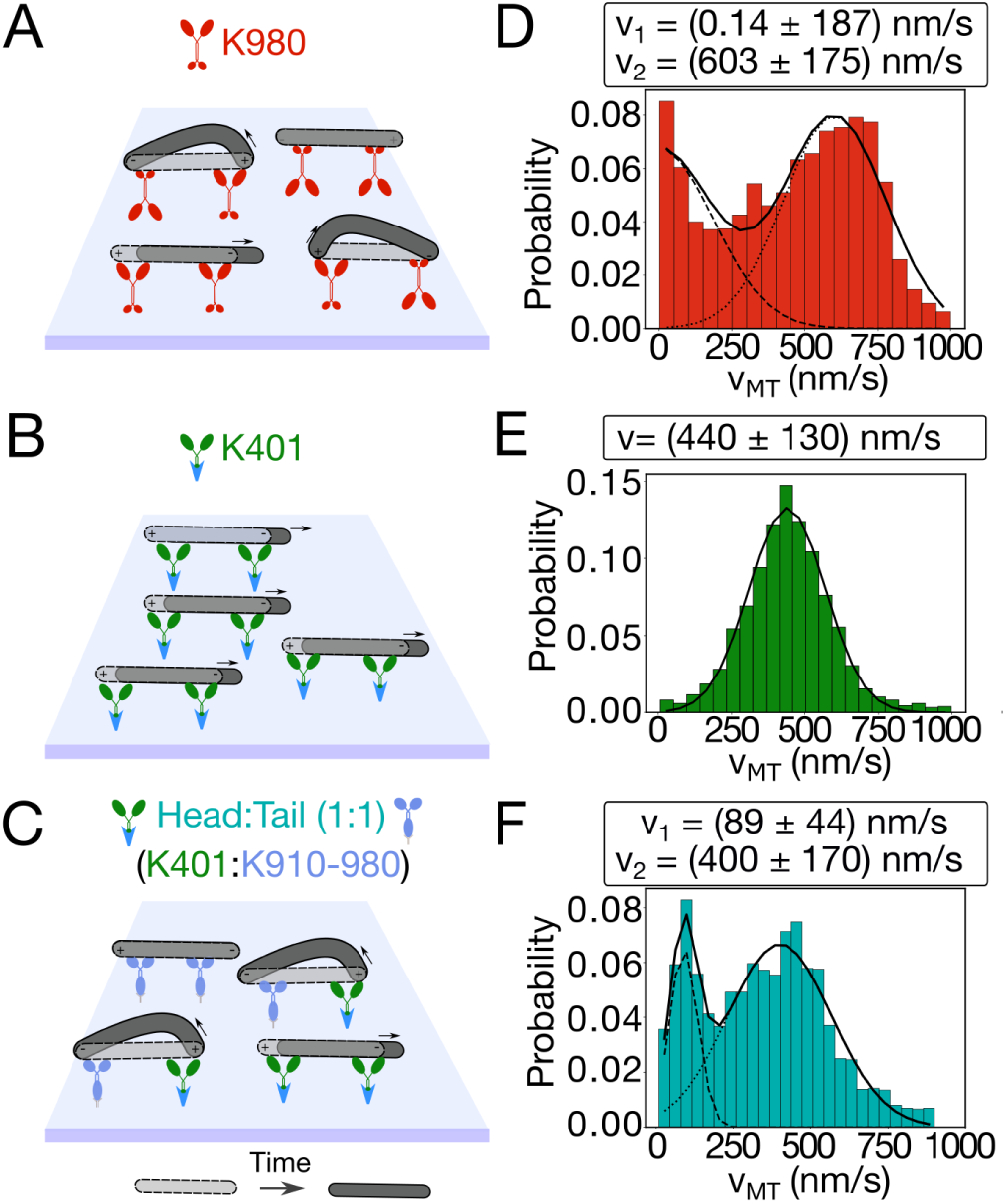
Equal head-tail mix reproduce bimodal velocity distribution of full-length kinesin-1. (A-C, left) Schematic of microtubule gliding assay for (A) full-length kinesin (K980), (B) truncated kinesin head (K401), (C) 1:1 head-tail mix (K401:K910-980). Microtubules bound at t = 0 (light gray), at t = t*_final_* (dark gray). (D-F) Corresponding microtubule gliding velocity probability distribution of V_MT_. The distributions for (D) K980 and (F) head:tail were fitted with a bimodal Gaussian (black line), as the sum of two single Gaussian components (dashed lines). (E) Unimodal Gaussian distribution was fitted to microtubule velocities driven by the minimal K401. (*N*_K980_ = 298, *N*_K401_ = 303, *N*_Head-Tail_ = 316)

To test this, we analyzed the velocity distribution of microtubules driven by K980 (Fig. 6A, D), just K401(Fig. 6B, E) and head-tail mix (K401:K910-980) (Fig. 6C, F). Both the K980 and motor-tail mix showed a bimodal velocity distribution. A bimodal Gaussian fit to the velocity distribution of K980 reveals peaks near 0 nm/s and 600 nm/s, and for the head-tail-driven motility peaks near 90 nm/s and 400 nm/s (Fig. 6D, F), reflecting a mixture of distinct motility states. The higher-velocity peaks likely corresponds to microtubules transported by active motor heads, while the near-zero peak may represent microtubules bound via the tail domain, which interacts with microtubules but does not generate movement. In contrast, a unimodal distribution was observed for gliding assays with only K401, fitted with a single peaked Gaussian function with peak velocity of 440 nm/s (Figs. 6E).

Thus, in addition to spatial patterns, an equal ratio of head and tail domains also reproduces the temporal dynamics observed with full-length kinesin. To further test this model, we next examined whether similar patterns persist when the ratio of tail domain and motor head is varied.

### Microtubule bending during gliding emerges from head-tail domain competition

To investigate how the balance between motor and tail domains affects microtubule velocity and spatial organization, we performed gliding assays across a range of head-to-tail ratios (*C*_Tail_*/C*_Tail+Head_, Fig. 7A). Increasing the proportion of K910-980 (*C*_Tail_) resulted in a shift in velocity distribution: in the total absence of the tail (*C*_Tail_*/C*_Tail+Head_ = 0) it resulted in a sharp single peak at high velocity, at a ratio of 0.25 a broad distribution of velocities was observed and eventually for a ratio of 0.5, two distinct peaks appeared near 0 nm/s and 400 nm/s, indicating coexisting stalled and motile microtubules. At higher ratios (0.75-1), velocities dropped down to ∼0 nm/s as tail binding dominated. Mean velocities declined sharply between ratios 0.5 and 0.75 with broader variability (Fig. 7B).

**Figure 7:**
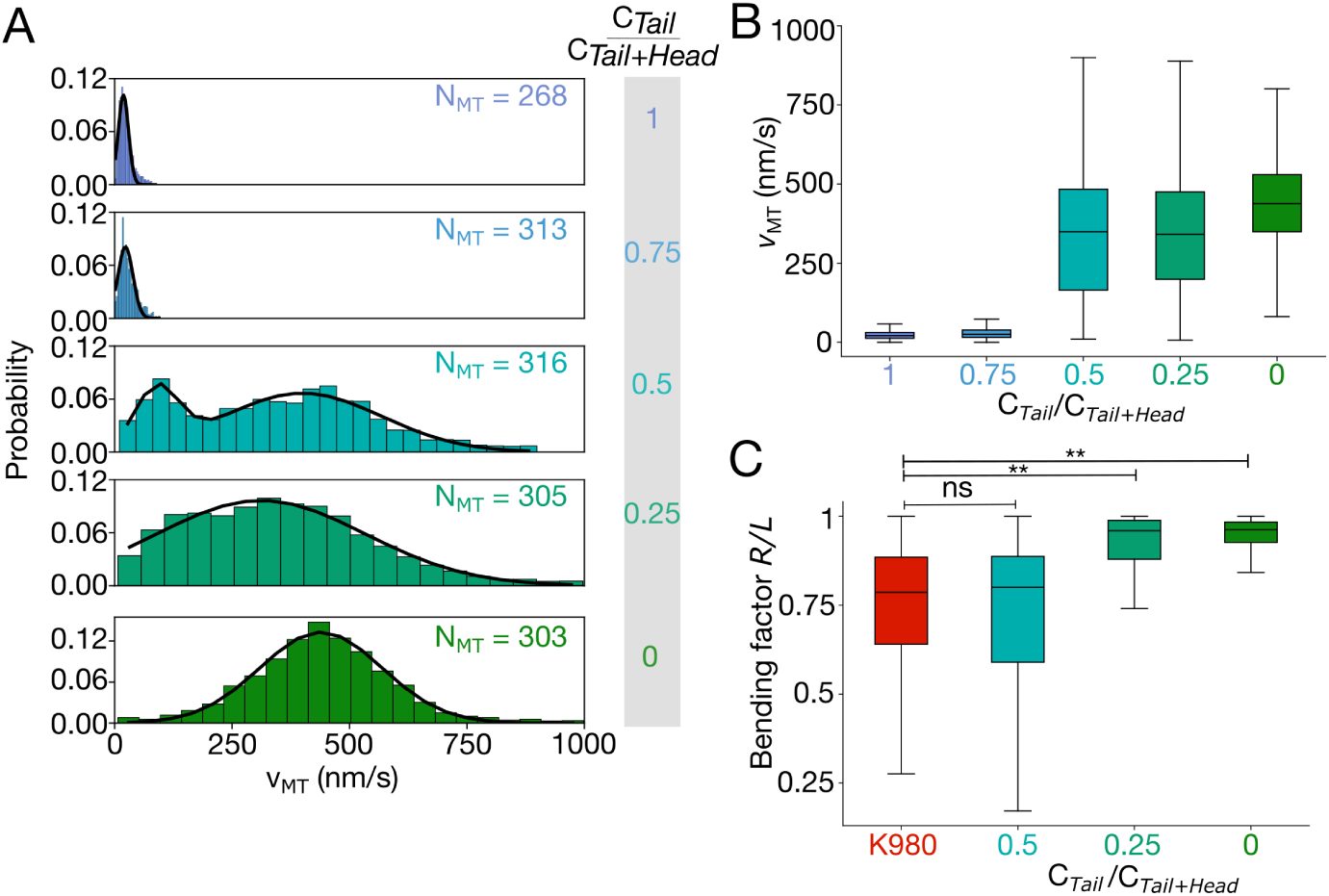
Head-tail ratio modulates microtubule gliding dynamics and curvature. (A) Microtubule velocity (*v*_MT_) distributions for varying *C*_Tail_*/C*_Tail+Head_ ratios, fitted with unimodal or bimodal Gaussians (Gauss fit sum of two, black lines). *C*_Tail_*/C*_Tail+Head_ = 1 corresponds to K910-980 and *C*_Tail_*/C*_Tail+Head_ = 0 corresponds to K401. (*N*_1_ = 268*, N*_0.75_ = 313*, N*_0.5_ = 316*, N*_0.25_ = 305*, N*_0_ = 303) (B) Box plot of mean microtubule velocity (*v̄*_MT_) as function of *C*_Tail_*/C*_Tail+Head_. Error bars: SD. (C) Bending factor (*R/L*) plotted for each ratio and compared to K980. ns: not statistical significant, ^∗∗^*p <* 0.01. (*N_K_*_980_ = 298*, N*_0.5_ = 316*, N*_0.25_ = 305*, N*_0_ = 303)

To assess spatial effects across head-tail mixtures, we analyzed the microtubule bending factor (*R/L*) at different ratios (Fig. 7C). As tail concentration increased, microtubule curvature also increased, with the lowest *R/L* values observed at 0.5 and 0.75, similar to full-length kinesin (K980). In contrast, at a tail fraction of 0.25 and 0.0, microtubules remained mostly straight (*R/L* ≈ 0.9), resembling K401-driven motility. Statistical comparison of bending factors showed no significant difference between the 1:1 head-tail mixture and full-length K980, while other conditions differed significantly. This suggests that significant bending and the associated spatial patterns only emerge when the tail domain is present in at least equimolar amounts.

Thus, while the tail domain slows microtubule gliding motility, collective spatial and temporal patterns emerge only at equimolar head-tail ratios.

## Discussion

In this study, we compare collective microtubule transport by a full-length Drosophila kinesin-1 (K980) to a truncated motor domain construct (K401). Unlike K401, which produced processive, straight-gliding microtubules, K980 generated striking length-dependent spatial patterns including bending, oscillations, and loops, along with stop-and-go motion. To explore the role of the C-terminal tail, we engineered a kinesin tail construct (residues 910-980) and confirmed that it binds microtubules but does not support gliding. By reconstituting equimolar mixtures of head and tail constructs, we reproduced the spatiotemporal patterns seen with K980, suggesting that the head and tail compete for microtubule engagement.

The emergence of spatial patterns in K980-driven gliding was particularly striking, as similar behaviors typically require artificial constraints such as pinning (Bourdieu et al., 1995) or clamping (Gittes et al., 1996; Vilfan et al., 2019; Yadav et al., 2024; Liu et al., 2011) of filament ends. In our assays no such constraints were added. This emphasizes the role of kinesin tails as anchors or pinning sites, when bound to microtubules, which can bind anywhere along the length of the filament. While K401-driven gliding showed occasional curvature in long filaments, nearly all K980-driven microtubules displayed dynamic patterns, supporting the interpretation of a mechanical competition between transport and immobilization. K980 also induced frequent pauses and velocity fluctuations, stop-and-go motion, resulting in a bimodal velocity distribution. Similar profiles have been attributed to mixtures of active and inactive motors (Korten et al., 2016; Scharrel et al., 2014), but these studies did not report spatial patterns. Our findings suggest that tail-bound motors act as static anchors, altering both the trajectory and velocity of microtubules. Our findings while based on *in vitro* reconstitution, could help develop a mechanistic explanation for previous observations of kinesin-dependent cytoplasmic flows seen in *Drosophila* embryos (Monteith et al., 2016). These studies had provided evidence pointing to the importance of kinesin-1 driven sliding (Lu et al., 2016) and a role for the heavy chain (Palacios and Johnston, 2002). Such models are generally based on a combination of filament sliding and mechanical feedback. However, a more recent model of self organizated hydro-dynamic flows arising from cargo transport by kinesin motors has been shown to reconcile well to experiment (Dutta et al., 2023). While this theory sheds new light on the emergence of cytoplasmic streaming *in vivo*, the mechanistic role of the kinesin-1 heavy chain, as the primary driver of such flows is still not clear. We believe our experiments could help expand the previous theory to build a complete picture from molecular mechanisms to embryo-scale flows.

Although some kinesin tails have been reported to exhibit diffusive motion (Furuta and Toyoshima, 2008; Shin et al., 2015), we find the kinesin-1 tail is static on binding to filaments. Individual tail proteins are seen to form dense puncta. In gliding assays, the tail does no without gliding or diffusion. This is consistent with prior observations in kinesin tail structures seen in fungi (Straube et al., 2006) and humans (Seeger et al., 2012). Despite similar binding densities to the motor domain, tail-bound microtubules exhibit distinct static behavior.

Our results support what we term the coin-toss model: in the absence of orientation control in immobilization, full-length kinesin binds the surface randomly, exposing either the motor or tail to microtubules This head-tail antagonism explains the observed spatial and temporal patterns. Importantly, this behavior was recapitulated by a 1:1 head-to-tail mixture, while deviating ratios disrupted pattern formation, further supporting the model.

In conclusion, our *in vitro* reconstitution demonstrates that a non-diffusive, microtubule-binding kinesin-1 tail can modulate microtubule transport when ori-entation is not controlled. This internal competition between head and tail domains provides a minimal mechanism for kinesin-mediated regulation of microtubule dynamics and patterning. Such antagonism may contribute to kinesin-1 roles in microtubule sliding, cytoplasmic streaming, and selective cargo transport *in vivo*.

## Methods

### Key Resources

File of key resources is provided with the submission.

### Experimental model details

All kinesin constructs were expressed and purified from *Escherichia coli* BL21(DE3) cells. *Escherichia coli* DH5*α* cells was used for transformation and cloning purposes.

### Method details

### Full-Length and truncated kinesin-1 Constructs

The *Drosophila* kinesin-1 full-length construct K980 (Addgene #129762, (Hancock and Howard, 1998)) and the truncated K401 construct (Addgene #15960, (Subramanian and Gelles, 2007)) were expressed in *E. coli* BL21(DE3) cells. K401 was expressed with 20 *µ*g/ml biotin for *in vivo* biotinylation (Berliner et al., 1994). Both constructs were purified using cobalt-affinity chromatography and analyzed by SDS-PAGE.

### Cloning and purification of tail and IAK-deleted constructs

The kinesin-1 tail domain (K910-980) was cloned by Gibson assembly using fragments from K980 and a GFP vector (Addgene #15219, (Woehlke et al., 1997)), verified by restriction digestion, and expressed in *E. coli* BL21(DE3). IAK-deleted mutants of the full-length construct (K936) were generated via PCR-based mutagenesis. both constructs were purified by cobalt-affinity chromatography and analyzed by SDS-PAGE.

### Rigor kinesin-1 mutant (K980(R210A)

The *Drosophila* kinesin-1 rigor mutant, K980(R210A) (Addgene #129763, a gift from William Hancock) was expressed in similar way as of K980 and purified using cobalt affinity chromatography with lysis buffer (50 mM sodium phosphate, 300 mM NaCl, 1 mM MgCl_2_, 0.1 mM MgATP, 5 mM *β*-mercaptoethanol and 0.1 mM PMSF, pH 8.0) supplemented with 150 mM imidazole (pH 7). The purified protein was analyzed by SDS-PAGE.

### GFP tagged truncated human kinesin-1, K560

A GFP-tagged human kinesin-1 truncated construct, K560 (Addgene #15219, (Woehlke et al., 1997)) was expressed in presence of 0.1 mM IPTG at 22^◦^C for 15 hours and purified using cobalt affinity chromatography.

All purified kinesin constructs were flash-frozen in 1 M sucrose and stored at −80^◦^C.

#### Purification and labelling of goat brain tubulin

Tubulin was purified from goat brain using a temperature-based polymerization-depolymerization method (Castoldi and Popov, 2003), with slight modifications as described previously (Jain et al., 2022; Basu et al., 2025). Purified tubulin was labeled with TAMRA using N-succinimidyl ester during a polymerization step, followed by one additional polymerization-depolymerization cycle (Hyman et al., 1991).

#### Reconstituting ***in vitro*** microtubule gliding assays

### Microtubule assembly

Taxol-stabilized microtubules were assembled by incubating 30 *µ*M goat tubulin with 5 *µ*M rhodamine-labeled tubulinand 10 % glycerol in PEM at 37 ^◦^C. Polymerized microtubules were stabilized with 20 *µ*M taxol (Paclitaxel; Cytoskeleton Inc.,Denver, CO, USA), pelleted at 150,000 g (TLA 100.3 rotor, Beckman Coulter), and resuspended in taxol-containing PEM.

### Gliding assay

Gliding assays were performed in flow chambers assembled from acetone-KOH washed 22 mm x 22 mm coverslips (No. 1.5, VWR; Avantor, USA) and 22 mm x 60 mm slides (HiMedia, India), separated by double-sided Kapton tape (Ted Pella Inc., USA). For each condition, 1 *µ*M protein was introduced into the chamber.

### Full-length kinesin-1 (K980)

K980 (1 *µ*M) were nonspecifically bound to the glass by direct incubation (15 min), followed by 1.5 mg/ml casein blocking.

### Truncated kinesin-1 motor domain construct (K401)

K401 (1 *µ*M) was specifically bound to streptavidin-coated glass surfaces by direct incubation (15 min), following pre-treatment with 1 mg/ml streptavidin and 1.5 mg/ml casein blocking. This ensured correct orientation with motor domains exposed to bind to micro-tubules.

### Kinesin mutant (K980(R210A))

The rigor kinesin mutant, K980(R210A) was used to determine the lower limit of detection of velocity in gliding assays. 66 *µ*g/ml of the purified protein were non-specifically bound to glass by direct incubation (15 min). 1.5 mg/ml casein was used for blocking the glass surface.

### Kinesin tail domain (K910-980)

GFP-tagged tail domains (1 *µ*M) were specifically immobilized on glass surfaces using a GFP nanobody (Addgene #61838, (Katoh et al., 2015)), ensuring that the tail domain was exposed for microtubule binding. The nanobody was purified from a pGEX6P1-GFP-nanobody construct as described previously (Yadav et al., 2024) (gift from Kazuhisa Nakayama, Addgene plasmid 61838, urlhttp://n2t.net/addgene:61838).

### IAK deletion mutant (K936)

1 *µ*M of purified K936 was non-specifically bound to the glass surface by direct incubation (15 min) and followed by 1.5 mg/ml casein for surface passivation.

### Different head-tail mixtures

The different molar ratios of head and tail were flowed into glass coverslips coated with 2 mg/ml of each of streptavidin and GFP nanobody. The total protein concentration was 1 *µ*M. *C*_Tail_*/C*_Tail+Head_ = 0.25: 0.75 *µ*M of K401 and 0.25 *µ*M of K910-980. *C*_Tail_*/C*_Tail+Head_ = 0.5: 0.5 *µ*M of K401 and 0.5 *µ*M of K910-980. *C*_Tail_*/C*_Tail+Head_ = 0.75: 0.25 *µ*M of K401 and 0.75 *µ*M of K910-980.

### Motor surface density

For all assays, kinesin constructs were introduced at a total concentration of 1 *µ*M, whether used individually or in combination. We expect an average motor density of ≈ 100 motors/*µ*m^2^ based on previous reports using similar concentrations (Scharrel et al., 2014; Monzon et al., 2019).

### Motility buffer

Rhodamine-labeled microtubules in PEM buffer containing 20 *µ*M paclitaxel were flowed in and incubated for 10 mins. Unbound microtubules were washed out by flowing PEM buffer containing 20 *µ*M paclitaxel, followed by motility buffer containing 5 mM MgCl_2_, Antifade mix: 1 *µ*M Glucose Oxidase, 1.5 *µ*M Catalase (all from SRL Pvt Ltd., Mumbai, India) and 50 mM Glucose, 1 % *β*-mercaptoethanol, 0.5 mg/ml casein and 1 mM MgATP in PEM.

#### Time-lapse fluorescence microscopy

Time-lapse imaging was performed using a Nikon TiE inverted epifluorescence microscope with a 60x NA 1.45 oil objective and a cooled Andor Clara2 CCD camera (Andor, Oxford Instruments) with a Halogen Lamp. The system was enclosed in a temperature-controlled chamber (Oko Labs, Italy) maintained at 37 ^◦^C. Images were acquired every 2-5 seconds for 5-10 minutes (700-900 ms exposure) using a TRITC filter for rhodamine-labeled microtubules and a FITC filter for GFP-tagged proteins.

#### Microtubule landing assay

Glass coverslips were coated with either 2 mg/ml of streptavidin or GFP nanobody, followed by 15 min incubation of either 1 *µ*M K401 or 1 *µ*M K910-980 respectively.

1.5 mg/ml casein was used for blocking. Rhodamine-labeled microtubules in PEM buffer containing 20 *µ*M paclitaxel was introduced into the flow chamber and incubated at 37*^o^*C for 10 mins. Unbound microtubules were then washed out by flowing PEM buffer containing 20 *µ*M paclitaxel. microtubules were then imaged using the TRITC filter using 100x (NA 1.4 oil) lens.

#### Microtubule motor co-localization assay

Microtubules were immobilized on acetone-KOH washed coverslips coated with rabbit anti-Tetramethylrhodamine antibody. GFP-tagged kinesin constructs (K560 or K910-980) were flowed into the flow chamber and incubated for 10-15 min at 37 *^o^*C. Imaging was performed with a 100x, NA 1.4 oil objective using TRITC (561 nm) for rhodamine-labeled microtubules and FITC (488 nm) for GFP. Channels were merged in FIJI to assess co-localization.

### Quantification and statistical analysis

#### Microtubule velocity analysis

Gliding assay time series were acquired at 0.5 fps for 100-120 s. The stacks were further background-subtracted and median-filtered in FIJI. Microtubule tips were tracked using FIESTA (Ruhnow et al., 2011) and instantaneous velocities were computed via 2*D* point-to-point displacement in FIESTA. Kymographs were generated using the Multi-Kymograph plugin in FIJI (Schindelin et al., 2012).

#### Bending factor (***R/L***)

For motile microtubules, contour length *L* and end-to-end distance R were measured interactively in FIJI using the line segment and measure tools. The ratio *R/L* was calculated for each microtubule and fitted against *L* using the flexural model described by Mizushima-Sugino (Mizushima-Sugano et al., 1983):

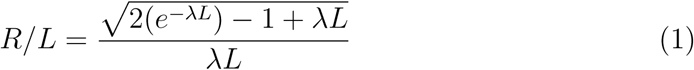

where *λ* is the bending stiffness parameter.

#### Quantifying microtubule patterns

Microtubule gliding behaviors were classified into three categories based on distinct spatial patterns observed in time-lapse image sequences. ”Bending” referred to microtubules displaying visible buckling or kinks along their contour during motility. ”Oscillation” described filaments that were anchored at one end and exhibited periodic, wave-like fluctuations, consistent with previously reported behaviors (Yadav et al., 2024). ”Looping” was defined as microtubules tethered at one end that formed circular or spiral-shaped profiles during movement. Only motile microtubule were included in the analysis. Microtubule patterns were visually analyzed, and each microtubule was assigned to the category that best described its overall behavior throughout the time-lapse sequence.

#### Microtubule density analysis

For both K401 and K910-980, microtubules were manually counted. Microtubule density for both heads and tails were further quantified by dividing the number of microtubules with the image area.

#### Statistical analysis

All data are presented as mean±standard deviation (SD), unless otherwise stated. Statistical comparisons between two groups were performed using unpaired two-tailed Student’s *t*-test. A p-value*>*0.05 was considered not significant (ns); significance levels are indicated as *p <* 0.05 (*), *p <* 0.01 (**). Data analysis was carried out in Python (v3.9.12) using NumPy (v1.21.5) (Harris et al., 2020) and SciPy (v1.7.3) (Virtanen et al., 2020). Data are based on at least three independent replicate experiments and include measurements from multiple fields of view per experiment.

## Acknowledgments

We are grateful to William Hancock for the full-length and rigor kinesin construct, K980 plasmid and Jeff Gelles for the K401-BCCP plasmid. We thank Ron Vale for the gift of the K560 construct.

## Author contributions

JB designed and performed the experiments, data analysis, made the figures, and wrote the first draft of the manuscript. KS performed control experiments and analyzed the data. AJ co-wrote the paper. CAA designed and supervised the research, obtained the funding and wrote and edited the article.

## Funding and additional information

JB is supported by a PhD studentship from IISER Pune. KS is supported by a fellowship from the Dept. of Biotechnology, Govt. of India (DBTHRDPMU/JRF/BET-23/I/2023-2024/356). The project was supported by IISER Pune faculty consumable funds.

## Data availability

This study includes no data deposited in external repositories

## Supplementary Information

1. Supplementary Table S1
2. Supplementary Figures S1 and S2
3. Supplementary Videos 1 to 6

## Supplementary Table

**Table S1:** Kinesin-1 constructs used in this study. Engineered and previously described plasmids are summarized; see Methods for details.

| Name | Full construct name | Source |
| --- | --- | --- |
| K980 | Drosophila Khc(1-980)-6xHis | Addgene plasmid no 129762, RRID:Addgene_129762 (Unpublished) |
| K401 | Drosophila Khc(1-401)-BCCP-6XHis | Addgene plasmid no 15960, RRID:Addgene_15960 (Subramanian and Gelles, 2007) |
| K560 | Human KIF5B(1-560)-GFP-6xHis | Addgene plasmid no 15219, RRID:Addgene_15219 (Woehlke et al., 1997) |
| K910-980 | Drosophila GFP-Khc(910-980)-6xHis | This study |
| K936 | Drosophila Khc(1-936)-6xHis | This study |
| K980(R210A) | Drosophila Khc(1-980, R210A) | Addgene plasmid no 129763, RRID:Addgene_129763 (Unpublished) |

## Supplementary Figures

**Figure S1:**
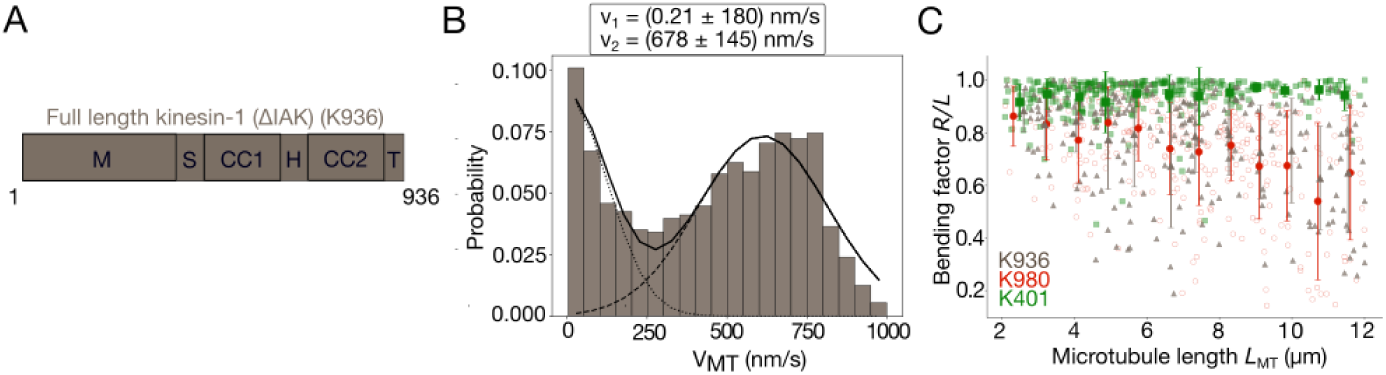
Collective microtubule transport driven by IAK deleted full-length kinesin (K936). (A) Schematic of the domain structure of full-length, kinesin-1 with IAK domain (944-946) deleted (ΔIAK), referred as K936. (B) Velocity probability distribution (*V*_MT_) measured from microtubule gliding driven by K936. The distribution was fitted to a bimodal Gaussian distribution (black line) showing peaks at ≈ 0 nm/s and ≈ 675 nm/s (N_K936_ = 224) (C) The bending factor (*R/L*) measured from K936 gliding assays as a function of microtubule length, L_MT_ (grey) compared to K980 (red) and K401 (green). Large symbols indicate binned means±SD (bin width: 1 *µ*m (N_K936_ = 224)

**Figure S2:**
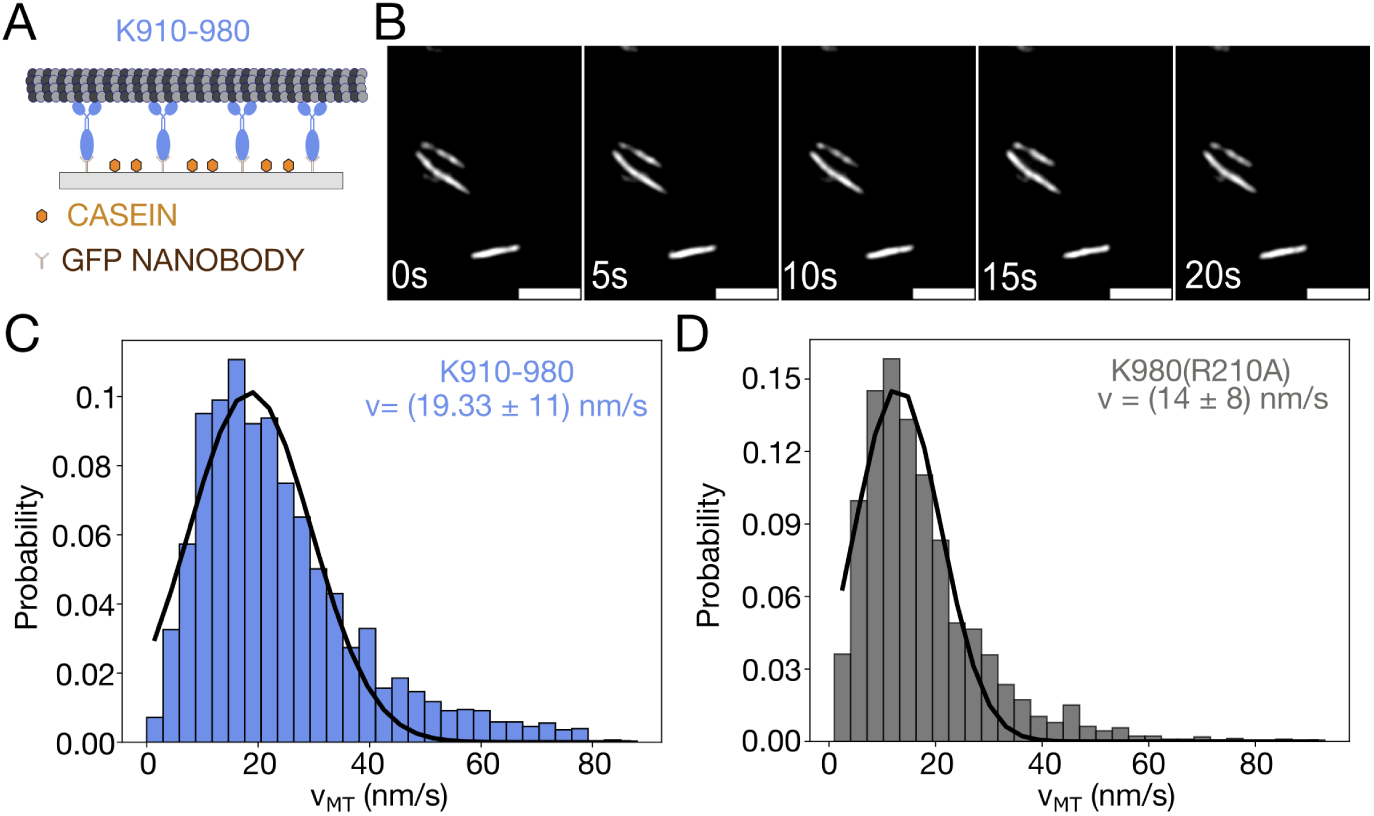
Kinesin-1 tail-bound microtubules are stationary in the presence of ATP. (A) Schematic of microtubule gliding assay with K910-980 immobilized on glass via GFP nanobody . (B) Time series of microtubules bound to K910-980. Scale bar: 5 *µ*m. (C,D) Instantaneous microtubule velocity distribution (VMT) for microtubules immobilized either on (C) K910-980 or (D) kinesin rigor mutant K980(R210A). Black lines: Gaussian fit. (*N*_K910-980_ = 268, *N*_K980(R210A)_ = 318)

## Supplementary Videos

**Supplementary Video 1.** Fluorescence time-lapse of rhodamine-labeled microtubules driven by full-length kinesin-1 (K980) in the presence of 1 mM ATP. Images acquired every 2 s using TRITC filter. Scale bar: 10 *µ*m. (mm:ss = minutes:seconds)

**Supplementary Video 2.** Fluorescence time-lapse of rhodamine-labeled microtubules driven by truncated kinesin head (K401) in the presence of 1 mM ATP. Images acquired every 2 s using TRITC filter. Scale bar: 10 *µ*m. (mm:ss = minutes:seconds)

**Supplementary Video 3.** Fluorescence time-lapse of rhodamine-labeled microtubules driven by IAK-deleted full-length kinesin-1 (K936) in the presence of 1 mM ATP. Images acquired every 2 s using TRITC filter. Scale bar: 10 *µ*m. (mm:ss = minutes:seconds)

**Supplementary Video 4.** Fluorescence time-lapse of rhodamine-labeled microtubules in presence of kinesin tail domain (K910-980) with 1 mM ATP. Images acquired every 1 s using TRITC filter. Scale bar: 2 *µ*m. (mm:ss = minutes:seconds)

**Supplementary Video 5.** Fluorescence time-lapse of rhodamine-labeled microtubules driven by a 1:1 mixture of K401 and K910-980 in the presence of 1 mM ATP. Images acquired every 2 s using TRITC filter. Scale bar: 10 *µ*m. (mm:ss = minutes:seconds)

**Supplementary Video 6.** Fluorescence time-lapse of rhodamine-labeled microtubules patterns in the presence of (A) K980 and (B) 1:1 mix of K401 and K910-980 with 1 mM ATP. Microtubules show bending (red), oscillation (magenta), looping (green), and gliding (blue). Images acquired every 2 s in TRITC filter. Scale bar: 5 *µ*m. (mm:ss = minutes:seconds)

## References

Basu, J., Soni, A., and Athale, C. A. (2025). Physical effects of crowdant size and concentration on collective microtubule polymerization. Biophysical Journal.

Berliner, E., Mahtani, H. K., Karki, S., Chu, L. F., Cronan, J. E., and Gelles, J. (1994). Microtubule movement by a biotinated kinesin bound to a streptavidin-coated surface. Journal of Biological Chemistry, 269(11):8610–8615.

Bieling, P., Telley, I. A., Piehler, J., and Surrey, T. (2008). Processive kinesins require loose mechanical coupling for efficient collective motility. EMBO reports, 9(11):1121– 1127.

Bourdieu, L., Duke, T., Elowitz, M., Winkelmann, D., Leibler, S., and Libchaber, A. (1995). Spiral defects in motility assays: a measure of motor protein force. Physical review letters, 75(1):176.

Brady, S. T. (1985). A novel brain atpase with properties expected for the fast axonal transport motor. Nature, 317(6032):73–5.

Castoldi, M. and Popov, A. V. (2003). Purification of brain tubulin through two cycles of polymerization-depolymerization in a high-molarity buffer. Protein Expr Purif, 32(1):83–8.

Crevenna, A. H., Madathil, S., Cohen, D. N., Wagenbach, M., Fahmy, K., and Howard, J. (2008). Secondary structure and compliance of a predicted flexible domain in kinesin-1 necessary for cooperation of motors. Biophysical journal, 95(11):5216–5227.

Dutta, S., Farhadifar, R., Lu, W., Kabacaoğlu, G., Blackwell, R., Stein, D. B., Lakonishok, M., Gelfand, V. I., Shvartsman, S. Y., and Shelley, M. J. (2023). Self-organized intracellular twisters. ArXiv, page arXiv:2304.02112v2.

Farrell, C. M., Mackey, A. T., Klumpp, L. M., and Gilbert, S. P. (2002). The role of atp hydrolysis for kinesin processivity. Journal of Biological Chemistry, 277(19):17079– 17087.

Furuta, K. and Toyoshima, Y. Y. (2008). Minus-end-directed motor ncd exhibits processive movement that is enhanced by microtubule bundling in vitro. Current Biology, 18(2):152–157.

Gittes, F., Meyhöfer, E., Baek, S., and Howard, J. (1996). Directional loading of the kinesin motor molecule as it buckles a microtubule. Biophysical journal, 70(1):418– 429.

Hancock, W. O. and Howard, J. (1998). Processivity of the motor protein kinesin requires two heads. The Journal of cell biology, 140(6):1395–1405.

Harris, C. R., Millman, K. J., Van Der Walt, S. J., Gommers, R., Virtanen, P., Cournapeau, D., Wieser, E., Taylor, J., Berg, S., Smith, N. J., et al. (2020). Array programming with numpy. nature, 585(7825):357–362.

Hirokawa, N., Noda, Y., Tanaka, Y., and Niwa, S. (2009). Kinesin superfamily motor proteins and intracellular transport. Nat. Rev. Mol. Cell Biol, 10:682–696.

Howard, J., Hudspeth, A., and Vale, R. D. (1989). Movement of microtubules by single kinesin molecules. Nature, 342(6246):154–158.

Hsu, H.-P., Paul, W., and Binder, K. (2010). Polymer chain stiffness vs. excluded volume: A monte carlo study of the crossover towards the worm-like chain model. Europhysics Letters, 92(2):28003.

Hyman, A., Drechsel, D., Kellogg, D., Salser, S., Sawin, K., Steffen, P., Wordeman, L., and Mitchison, T. (1991). [39] preparation of modified tubulins. In Methods in enzymology, volume 196, pages 478–485. Elsevier.

Jain, K., Basu, J., Roy, M., Yadav, J., Patil, S., and Athale, C. A. (2022). Polymerization kinetics of tubulin from mung seedlings modeled as a competition between nucleation and GTP-hydrolysis rates. Cytoskeleton, 78(9):436–447.

Jolly, A. L., Kim, H., Srinivasan, D., Lakonishok, M., Larson, A. G., and Gelfand, V. I. (2010). Kinesin-1 heavy chain mediates microtubule sliding to drive changes in cell shape. Proceedings of the National Academy of Sciences, 107(27):12151–12156.

Jon Kull, F., Sablin, E. P., Lau, R., Fletterick, R. J., and Vale, R. D. (1996). Crystal structure of the kinesin motor domain reveals a structural similarity to myosin. Nature, 380(6574):550–555.

Kaan, H. Y. K., Hackney, D. D., and Kozielski, F. (2011). The structure of the kinesin-1 motor-tail complex reveals the mechanism of autoinhibition. Science, 333(6044):883– 885.

Katoh, Y., Nozaki, S., Hartanto, D., Miyano, R., and Nakayama, K. (2015). Architectures of multisubunit complexes revealed by a visible immunoprecipitation assay using fluorescent fusion proteins. Journal of cell science, 128(12):2351–2362.

Korten, T., Chaudhuri, S., Tavkin, E., Braun, M., and Diez, S. (2016). Kinesin-1 Expressed in Insect Cells Improves Microtubule in Vitro Gliding Performance, Long-Term Stability and Guiding Efficiency in Nanostructures. IEEE Transactions on Nanobioscience, 15(1):62–69.

Kratky, O. and Porod, G. (1949). X-ray investigiation of chain molecules in solution. Recl. Trav. Chim. Pays Bas, 68:1106–1122.

Liu, L., Tüzel, E., and Ross, J. L. (2011). Loop formation of microtubules during gliding at high density. Journal of Physics: Condensed Matter, 23(37):374104.

Lu, W., Winding, M., Lakonishok, M., Wildonger, J., and Gelfand, V. (2016). Microtubule-microtubule sliding by kinesin-1 is essential for normal cytoplasmic streaming in drosophila oocytes. Proc. Natl. Acad. Sci. U. S. A, 113:E4995–E5004.

Mickolajczyk, K. J., Deffenbaugh, N. C., Ortega Arroyo, J., Andrecka, J., Kukura, P., and Hancock, W. O. (2015). Kinetics of nucleotide-dependent structural transitions in the kinesin-1 hydrolysis cycle. Proceedings of the National Academy of Sciences, 112(52):E7186–E7193.

Mizushima-Sugano, J., Maeda, T., and Miki-Noumura, T. (1983). Flexural rigidity of singlet microtubules estimated from statistical analysis of their contour lengths and end-to-end distances. Biochimica et Biophysica Acta (BBA)-General Subjects, 755(2):257–262.

Monteith, C., Brunner, M., Djagaeva, I., Bielecki, A., Deutsch, J., and Saxton, W. (2016). A mechanism for cytoplasmic streaming: Kinesin-driven alignment of microtubules and fast fluid flows. Biophys. J, 110:2053–2065.

Monzon, G. A., Scharrel, L., Santen, L., and Diez, S. (2019). Activation of mammalian cytoplasmic dynein in multimotor motility assays. Journal of cell science, 132(4):jcs220079.

Moua, P., Fullerton, D., Serbus, L. R., Warrior, R., and Saxton, W. M. (2011). Kinesin-1 tail autoregulation and microtubule-binding regions function in saltatory transport but not ooplasmic streaming. Development, 138(6):1087–1092.

Navone, F., Niclas, J., Hom-Booher, N., Sparks, L., Bernstein, H. D., McCaffrey, G., and Vale, R. D. (1992). Cloning and expression of a human kinesin heavy chain gene: interaction of the cooh-terminal domain with cytoplasmic microtubules in transfected cv-1 cells. The Journal of cell biology, 117(6):1263–1275.

Palacios, I. and Johnston, D. (2002). Kinesin light chain-independent function of the kinesin heavy chain in cytoplasmic streaming and posterior localisation in the drosophila oocyte. Development, 129:5473–5485.

Ruhnow, F., Zwicker, D., and Diez, S. (2011). Tracking single particles and elongated filaments with nanometer precision. Biophysical journal, 100(11):2820–2828.

Scharrel, L., Ma, R., Schneider, R., Jülicher, F., and Diez, S. (2014). Multimotor transport in a system of active and inactive kinesin-1 motors. Biophysical journal, 107(2):365–372.

Schindelin, J., Arganda-Carreras, I., Frise, E., Kaynig, V., Longair, M., Pietzsch, T., Preibisch, S., Rueden, C., Saalfeld, S., Schmid, B., et al. (2012). Fiji: an open-source platform for biological-image analysis. Nature methods, 9(7):676–682.

Scholey, J. M., Porter, M. E., Grissom, P. M., and McIntosh, J. R. (1985). Identification of kinesin in sea urchin eggs, and evidence for its localization in the mitotic spindle. Nature, 318(6045):483–6.

Seeger, M. A. and Rice, S. E. (2010). Microtubule-associated protein-like binding of the kinesin-1 tail to microtubules. Journal of Biological Chemistry, 285(11):8155–8162.

Seeger, M. A., Zhang, Y., and Rice, S. E. (2012). Kinesin tail domains are intrinsically disordered. Proteins: Structure, Function, and Bioinformatics, 80(10):2437–2446.

Shin, Y., Du, Y., Collier, S. E., Ohi, M. D., Lang, M. J., and Ohi, R. (2015). Biased brownian motion as a mechanism to facilitate nanometer-scale exploration of the microtubule plus end by a kinesin-8. Proceedings of the National Academy of Sciences, 112(29):E3826–E3835.

Shubeita, G. T., Tran, S. L., Xu, J., Vershinin, M., Cermelli, S., Cotton, S. L., Welte, M. A., and Gross, S. P. (2008). Consequences of motor copy number on the intracellular transport of kinesin-1-driven lipid droplets. Cell, 135(6):1098–1107.

Straube, A., Hause, G., Fink, G., and Steinberg, G. (2006). Conventional kinesin mediates microtubule-microtubule interactions in vivo. Molecular biology of the cell, 17(2):907–916.

Subramanian, R. and Gelles, J. (2007). Two distinct modes of processive kinesin movement in mixtures of atp and amp-pnp. The Journal of general physiology, 130(5):445– 455.

Vale, R. D., Reese, T. S., and Sheetz, M. P. (1985). Identification of a novel force-generating protein, kinesin, involved in microtubule-based motility. Cell, 42(1):39–50.

Vilfan, A., Subramani, S., Bodenschatz, E., Golestanian, R., and Guido, I. (2019). Flagella-like beating of a single microtubule. Nano letters, 19(5):3359–3363.

Virtanen, P., Gommers, R., Oliphant, T. E., Haberland, M., Reddy, T., Cournapeau, D., Burovski, E., Peterson, P., Weckesser, W., Bright, J., et al. (2020). SciPy 1.0: Fundamental Algorithms for Scientific Computing in Python. Nature Methods, 17:261–272.

Visscher, K., Schnitzer, M. J., and Block, S. M. (1999). Single kinesin molecules studied with a molecular force clamp. Nature, 400(6740):184–189.

Winding, M., Kelliher, M. T., Lu, W., Wildonger, J., and Gelfand, V. I. (2016). Role of kinesin-1–based microtubule sliding in drosophila nervous system development. Proceedings of the National Academy of Sciences, 113(34):E4985–E4994.

Woehlke, G., Ruby, A. K., Hart, C. L., Ly, B., Hom-Booher, N., and Vale, R. D. (1997). Microtubule interaction site of the kinesin motor. Cell, 90(2):207–216.

Yadav, S. A., Khatri, D., Soni, A., Khetan, N., and Athale, C. A. (2024). Wave-like oscillations of clamped microtubules driven by collective dynein transport. Biophysj, 123(4):509–524.

Yildiz, A., Tomishige, M., Vale, R. D., and Selvin, P. R. (2004). Kinesin walks hand-over-hand. Science, 303(5658):676–678.

